# Loss of CaMKI function disrupts salt aversive learning in *C. elegans*

**DOI:** 10.1101/149682

**Authors:** Jana P. Lim, Holger Fehlauer, Dominique A. Glauser, Anne Brunet, Miriam B. Goodman

## Abstract

The ability to adapt behavior to environmental fluctuations is critical for survival of organisms ranging from invertebrates to mammals. *Caenorhabditis elegans* can learn to avoid sodium chloride when it is paired with starvation. This behavior is likely advantageous to avoid areas without food. While some genes have been implicated in this salt aversive learning behavior, critical genetic components, and the neural circuit in which they act, remain elusive. Here, we show that the sole worm ortholog of mammalian CaMKI/IV, CMK-1, is essential for salt aversive learning behavior in *C. elegans*. We find that CMK-1 acts in the primary salt-sensing ASE neurons to regulate this behavior. By characterizing the intracellular calcium dynamics in ASE neurons using microfluidics, we find that loss of *cmk-1* leads to an altered pattern of sensory-evoked calcium responses that may underlie salt aversive learning. Our study implicates the conserved CaMKI/CMK-1 as an essential cell-autonomous regulator for behavioral plasticity to environmental salt in *C. elegans*.

**Significance Statement:** Like other animals, the nematode *Caenorhabditis elegans* depends on salt for survival and navigates toward high concentrations of this essential mineral. Because salt generates osmotic stress at high concentrations, it also threatens the survival of small terrestrial animals like *C. elegans*. A growing body of evidence indicates that *C. elegans* balances these factors through a process called salt aversive learning. We show that this behavior depends on expression of a calcium/calmodulin-dependent kinase, CMK-1, in the ASE salt sensing neurons and that salt-induced calcium signals in the ASE neurons are less sensitive to salt conditioning in animals lacking CMK-1 function. Our study identifies CMK-1 and sensory neurons as key factors in this form of behavioral plasticity.

## Introduction

Behavioral plasticity allows animals to survive a constantly changing environment. This trait enables organisms to seek environments rich in essential resources like food and potential mates, and to avoid barren or perilous ones. Commensurate with its importance, there have been widespread efforts to identify the genes and neural circuits involved in behavioral plasticity. The nematode *Caenorhabditis elegans* exhibits behavioral plasticity in response to a variety of external stimuli, including temperature, touch, smells, and tastes (Ardiel and Rankin, 2010). For this reason and because of its well-characterized nervous system and ease of genetic manipulation, *C. elegans* is as an informative model system for investigating behavioral plasticity.

All animals require salt (sodium chloride) for survival and, as a result, salt-seeking is a highly conserved animal behavior (Geerling and Loewy, 2008; Hurley and Johnson, 2015). However, high salt can also be a threat especially to small, terrestrial animals like *C. elegans* because it generates osmotic stress (Choe, 2013; Kim and Jin, 2015). Thus, the worm experiences intense pressures to balance the need for salt with the need to avoid high concentrations. A considerable body of evidence establishes that *C. elegans* nematodes seek salt by performing positive chemotaxis (Ward, 1973; Bargmann and Horvitz, 1991) and this attraction can be transformed into aversion (negative chemotaxis) following conditioning in solutions rich in salt and poor in bacterial food (Saeki et al., 2001; Tomioka et al., 2006). Although several signaling pathways (Table 1), including those linked to insulin signaling (Tomioka et al., 2006; Vellai et al., 2006; Adachi et al., 2010; Oda et al., 2011; Kunitomo et al., 2013) modulate this salt aversive learning behavior, the majority of mutants analyzed thus far exhibit only modest deficits in learning ability. Thus, genes essential for salt aversive learning behavior and their cellular locus of action remain under-studied.

**Table 1.** Pathways implicated in salt aversive learning

| **Pathway** | **Citations** |
| --- | --- |
| G protein signaling | (Jansen et al., 2002; Hukema et al., 2006; Matsuki et al., 2006; Sakashita et al., 2008; Adachi et al., 2010; Kunitomo et al., 2013) |
| Insulin signaling | (Tomioka et al., 2006; Vellai et al., 2006; Adachi et al., 2010; Oda et al., 2011; Kunitomo et al., 2013) |
| Synaptic proteins | (Ishihara et al., 2002; Ikeda et al., 2008; Stetak et al., 2009; Iwata et al., 2011) |
| Neurotransmitters | (Hukema et al., 2008; Kano et al., 2008; Stetak et al., 2009) |
| Other | (Saeki et al., 2001; Iino and Yoshida, 2009; Sakai et al., 2013) |

Calcium and calmodulin (CaM)-dependent kinases play evolutionarily conserved roles in the nervous system including signal transduction, synaptic development, and behavioral plasticity. Among other functions, CaM kinases respond to changes in cytosolic calcium levels and regulate gene transcription in the nucleus. Although CaMKII is the most intensively studied, CaMKI and CaMKIV have been found to be important for specific functions such as developmental and activity-dependent growth (Hook and Means, 2001; Wayman et al., 2008; Cohen et al., 2015). Recently, the sole worm ortholog of mammalian CaMKI/IV, CMK-1, has emerged as a versatile regulator of neuronal plasticity in *C. elegans*. CMK-1 is expressed broadly throughout the worm’s nervous system (Kimura et al., 2002; Satterlee et al., 2004), and regulates behavioral plasticity in responses to thermal stimuli (Satterlee et al., 2004; Schild et al., 2014; Yu et al., 2014; Kobayashi et al., 2016). It also regulates developmental plasticity by affecting entry into the “dauer” diapause state in response to food deprivation (Neal et al., 2015), and synaptic plasticity by regulating expression of an AMPA-type glutamate receptor (Moss et al., 2016). However, whether CMK-1 is implicated in salt attraction or its transformation to salt aversion following conditioning is unknown.

Here, we show that CMK-1 is dispensable for salt attraction, but absolutely required for salt aversive learning behavior. Disrupting the only other CaM kinase encoded by the *C. elegans* genome, UNC-43 CaMKII, or other kinases AMP kinase and protein kinase C, has no detectable effect on either positive chemotaxis or its transformation to negative chemotaxis following conditioning in high-salt in the absence of bacterial food. By expressing wild-type CMK-1 in a neuron-specific manner, we find that *cmk-1* functions in the ASEL and ASER chemosensory neurons known to be required for chemotaxis (Bargmann and Horvitz, 1991). Having identified the ASE neurons as a critical cellular locus for *cmk-1* action, we analyzed salt-evoked calcium signals in wild-type and *cmk-1* mutant ASE neurons. We show that conditioning sufficient to induce salt aversive learning blunts salt-evoked calcium signals in both ASE dendrites. Finally, we observe that *cmk-1* loss results in aberrant calcium responses of these neurons. Our results indicate that CMK-1 is essential for behavioral plasticity in *C. elegans* by acting cell-autonomously in a specific neuronal circuit.

## Materials and Methods

### Worm strains and husbandry

*C. elegans* strains were cultured using standard conditions at 20°C on NGM plates seeded with OP50-1 bacteria. All experiments were performed using adult hermaphrodites. Transgenic worms were generated using microinjection, resulting in formation of extra-chromosomal arrays. A subset of the *C. elegans* strains we used were generated specifically for this study, the remainder were obtained from our in-house collections, as gifts from colleagues, or from the Caenorhabditis Genetics Center. Table 2 lists all strains, their origin, and the figures showing data from each strain.

**Table 2.** List of *C. elegans* strains used in this study

| Strain name | Genotype | Origin or citation | Figure |
| --- | --- | --- | --- |
| N2 | wild type, Bristol | CGC | Throughout |
| RB754 | *aak-2(ok524) X* | CGC | Fig 1C |
| RB781 | *pkc-1(ok563) V* | CGC | Fig 1C |
| VC127 | *pkc-2(ok328) X* | CGC | Fig 1C |
| VC1052 | *unc-43(gk452) IV* | CGC | Fig 1C |
| JT200 | *unc-43(sa200) IV* | CGC | Fig 1B |
| VC220 | *cmk-1(ok287) IV* | CGC | Throughout |
| PY1589 | *cmk-1(oy21) IV* | CGC | Fig 1C, 3B |
| VC691 | *ckk-1(ok1033) III* | CGC | Fig 2A |
| YT17 | *crh-1(tz2) III* | CGC | Fig 2D |
| MT9973 | *crh-1(n3315) III* | (Kauffman et al., 2010) | Fig 2D |
| PS3551 | *hsf-1(sy441) I* | CGC | Fig 2D |
| GN363 | *cmk-1(pg58) IV* | (Schild et al., 2014) | Fig 4D |
| GN634 | *cmk-1(ok287) IV; oyIs17[gcy-8p::GFP] V; pgEx167[cmk-1p::cmk-1(WT)::GFP; myo-3p::mCherry]* | This study | Fig 1D, 4A |
| DAG307 | *cmk-1(ok287) IV; domEx307[cmk-1p::cmk-1(1-348)T179A; unc-122p::RFP]* | (Schild et al., 2014) | Fig 2B |
| GN720 | *cmk-1(ok287) IV; domEx328[cmk-1p::cmk-1(K52A)::GFP; unc-122p::RFP]* | This study | Fig 2C |
| DAG610-3 | *domEx610-3[flp-6p::cmk-1::GFP; cmk-1p::QF; QUAS::cmk-1::mCherry; unc-122p::RFP]* | This study | Fig 3A |
| GN636 | *cmk-1(ok287) IV; domEx122[mec-3p::cmk-1(WT)::GFP; unc-122::RFP]* | This study | Fig 3B |
| GN640 | *cmk-1(ok287) IV; oyIs17[gcy-8p::GFP] V; pgEx171[flp-6p::cmk-1(WT)::GFP; myo-3p::mCherry]* | This study | Fig 3B |
| PY8395 | *cmk-1(oy21) IV; Ex[ceh-36Dp::cmk-1(WT); unc-122p::dsRed] line 1* | (Neal et al., 2015) | Fig 3B |
| DAG176 | *cmk-1(ok287) IV; domEx176[cmk-1p::cmk-1(WT)::NES::GFP; unc-122p::RFP]* | (Schild et al., 2014) | Fig 4B, C |
| DAG178 | *cmk-1(ok287) IV; domEx178[cmk-1p::cmk-1(WT)::GFP::NLS; unc-122p::RFP]* | (Schild et al., 2014) | Fig 4B, C |
| ZC1600 | *yxEx738 [flp-6p::GCaMP3; unc-122p::GFP]* | (Luo et al., 2014) | Fig 5, 6A – B, E, 7A, C, Extended data A |
| GN637, | *cmk-1(ok287) IV; yxEx738 [flp-* | This study | Fig 6C – E, 7B – |
| GN670 | *6p::GCaMP3; unc-122p::GFP* |  | C, Extended data<br>B |

### Salt conditioning

Following a previously reported method (Tomioka et al., 2006), we transferred adult hermaphrodites from culture plates to 1.5 ml microcentrifuge tubes containing one of two conditioning buffers prior to each assay. The first buffer contained no added NaCl (0 mM) and the second buffer contained 20 mM NaCl. Both buffers also contained (in mM): KPO_4_ (5, pH 6.0), MgSO_4_ and CaCl_2_ (1). We removed residual bacteria by washing animals once in fresh buffer and conditioned worms for the indicated time periods (between 5 minutes and 10 hours).

### Chemotaxis assays

We adapted methods for assessing salt chemotaxis and salt aversive learning from Tomioka *et al.* (Tomioka et al., 2006) and diagram the process in Figure 1A. We generated radial salt gradients on assay plates by placing a 5 mm thick high salt agar plug 5 mm away from the edge of the 3.5 cm assay plate (with a 5 mm thick agar plate) and leaving it in place for 3 hours at 23°C. The high salt, 2% agar plug contained (in mM): KPO_4_ (5, pH 6.0), MgSO_4_ (1), CaCl_2_ (1), NaCl (100). The assay 2% agar assay plate contained (in mM): KPO_4_ (5, pH 6.0), MgSO_4_ (1), CaCl_2_ (1). Immediately prior to the start of the chemotaxis assay, the plug was removed and a drop (1*μ*L) of sodium azide (0.1 M) was spotted on opposite sides of the assay plate to immobilize animals near their final positions.

**Figure 1.**
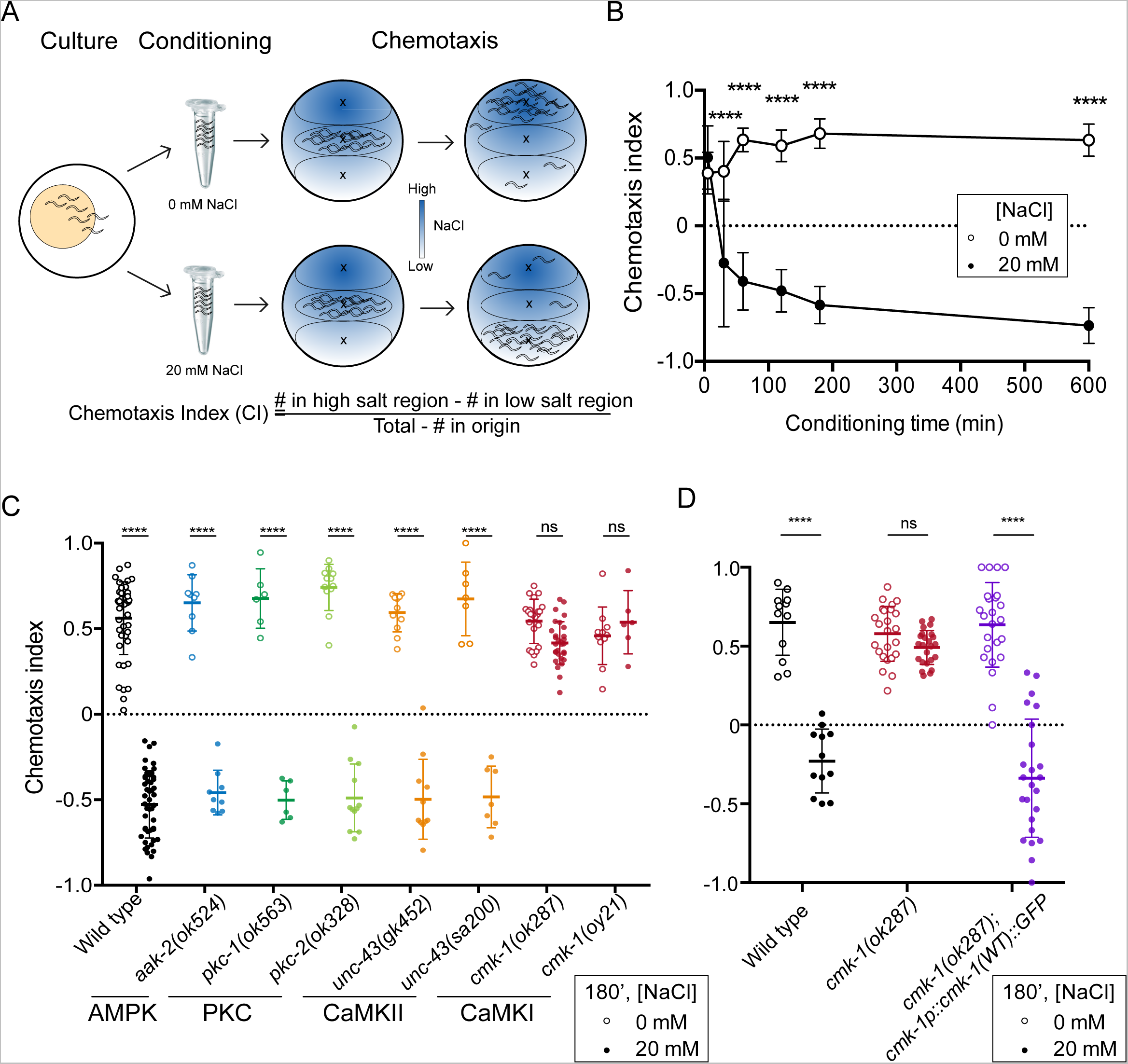
CMK-1 is required for salt chemotaxis learning. **A.** Procedure for testing salt aversive learning. (Left to right): Young adult (Day 1) worms are collected from culture plates, conditioned in food-free buffer containing 0 or 20 mM NaCl for 180 minutes, unless otherwise specified, transferred to the center of an assay plate pre-treated to create a radial salt gradient, allowed to migrate for 30 minutes. Results were quantified by computing a Chemotaxis Index (CI) (see Materials and Methods). **B.** Salt aversion development following several hours of conditioning in wild-type (N2) adults. Symbols are the mean ± SD of 3-9 assays for animals conditioned in 0 mM NaCl (open circles) or 20 mM NaCl (filled circles) for the indicated times. Data were pooled from 4 independent experiments consisting of 3-6 assays/condition. **C.** Chemotaxis behavior of kinase loss-of-function mutants conditioned without food and either 0 mM or 20 mM NaCl for 180 minutes (180’). Each circle is the result of a single assay; horizontal bars indicate the mean ± SD of all the data. Data were pooled from at least 2 independent experiments consisting of 3-6 replicates/condition. All assays were performed blind to genotype. A two-way ANOVA revealed a significant interaction between genotype and condition [*F*(7,237) = 62.34,*p* < 0.0001]. Sidak’s multiple comparisons test was used to assess the effect of conditioning for each genotype, which was significant at a level of *p* < 0.0001 (****) for all genotypes except for the two *cmk-1* alleles (ns). **D**. Transgenic expression of wild-type CMK-1::GFP under its own promoter restores wild-type salt aversive learning to *cmk-1(ok287)* mutants. Each symbol represents the results of a single assay; horizontal bars indicate mean ± SD of all assays. Data were pooled from at least 2 independent experiments consisting of 3-6 assays/condition. A two-way ANOVA revealed a significant interaction between genotype and condition [*F*(2, 114) = 44.2, *p* < 0.0001] and Sidak’s multiple comparisons test revealed a significant effect of conditioning (p < 0.0001, ****) for all cases except for *cmk-1(ok287)* (ns).

Conditioned worms were deposited in the middle of a 3.5 cm diameter radial salt gradient assay plate. Residual buffer was wicked off with a Kimwipe, and worms were allowed to crawl for 30 minutes at 20°C. The number of worms in each region (origin, low salt, high salt) was counted and used to compute a chemotaxis index:

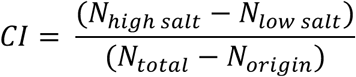

where *CI* is the chemotaxis and *N* is the number of worms in the indicated category. For each assay, we collected well-fed worms from a single growth plate (6 cm) into one 1.5 ml microcentrifuge tube for conditioning and then divided this sample into three equal aliquots to perform chemotaxis assays in triplicate.

### Locomotion assay

We collected adult (age: 1 day) worms from growth plates and conditioned them for 180 minutes, as described above for chemotaxis assays. Locomotion speed was assessed in the absence of food on standard NGM plates and we allowed animals to acclimate for 5 minutes before imaging. We collected short (1 minute) movies at 6x magnification at 20 frames/s (50 ms/frame) with a CCD camera (Mightex, M/N SME-B050-U) mounted on zoom lens (Navitar) with a C-mount adaptor and 0.5× lens and an illumination ring positioned in the same plane as the agar plate and providing radial illumination around it. Tracks showing locomotion were generated using the automated Parallel Worm Tracker software (Ramot et al., 2008) and were used to compute the average speed/track. The average speed of each sample was determined by computing the ensemble average of speed/track.

### Co-localization imaging

Co-labeling imaging for *flp-6p* and *cmk-1p* fluorescent reporters was performed on an Axioplan 2 Zeiss epifluorescence microscope equipped with an Axiocam camera and a 40x objective (air, NA = 0.95), as previously described (Hostettler et al., 2017).

### Subcellular localization imaging

We collected adult (age: 1 day) worms from growth plates and conditioned them for 180 minutes. Adult (age: 1 day) worms were conditioned in buffer containing 5 mM KPO_4_ (pH 6.0),1 mM MgSO_4_, 1 mM CaCl_2_, and 0 mM NaCl, 20 mM NaCl, or 0 mM NaCl with OP50-1 bacteria for 0, 10, 30, and 180 minutes. After conditioning we put the animals into a microfluidic chip (Nekimken et al., 2017) to restrict the worm’s motility during imaging. We took images at a Leica DMI 4000 B microscopy system, consisting of a cyan LED (Spectra X light engine, Lumencor), a fluorescence cube with a beam splitter (Chroma, Q495lp) and an emission filter (Chroma, HQ500lp), a 63x / 1.32 NA oil objective (Leica), and a Hamamatsu Orca-Flash 4.0LT digital CMOS camera. The subcellular localization was quantified by comparing the relative average fluorescence intensity of a hand-drawn ROI covering the nucleus versus the average fluorescence intensity of an ROI covering the cytoplasmic compartments each subtracted by the average fluorescence of an ROI outside the cell.

### *in vivo* calcium imaging

We collected worms (age: 1 day) expressing the *flp-6p:: GCaMP3* transgene and conditioned them for at least 180 minutes, using the same procedures and buffers as described for chemotaxis assays. Following conditioning, worms were positioned in a polydimethylsiloxane (PDMS) microfluidic device (MicroKosmos, LLC) that immobilizes the worm’s body while leaving the nose exposed to laminar buffer flow. The chip design is based on a device reported by Chronis *et al*. (Chronis et al., 2007). Under baseline conditions, the worm’s nose was exposed to a NaCl-free buffer. Chemosensory stimulation and imaging occurred during 90-second time blocks consisting of a control period (10 s), a 40-second NaCl pulse (20 mM), and a recovery period (40 s). The solutions were switched as described in Chalasani *et al*. (Chalasani et al., 2007) by a three-way valve (778360, The Lee Company) that was controlled by a power supply (6212B, Hewlett Packard). We added fluorescein (~20 mM) to the solution coming from the two side channels. The stream from these two channels is used to redirect the flow of the NaCl-free and the 20mM NaCl buffer and to visualize solution switches.

Calcium imaging of neurons of the worms in the chip was performed on a Leica DMI 4000 B equipped for epifluorescence imaging based on a cyan LED light source (Spectra X light engine, Lumencor), image splitting optics (W-view Gemini, Hamamatsu), and an sCMOS camera (Orca-Flash 4.0LT, Hamamatsu). We mounted a long-pass filter (Chroma, Q495lp) and an emission filter (Chroma, HQ500lp) in the beam splitter to collect GCaMP3 fluorescence. Fluorescence signals were collected using a 20x/0.4 NA objective (506200, Leica) and recorded at 10 Hz for the entire 90 s of the stimulus protocol.

All image sequences were analyzed using custom code running inside Fiji (Schindelin et al., 2012). Briefly, a desired region of the cell, the sensory ending (dendrite), cell body, or axons was selected manually in the first image of each sequence. In the first image the program identifies the brightest (*b*) and the dimmest (*d*) pixel in an area (5 x 5 *μ*m for cell bodies and 1 x 1 *μ*m for dendrites and axons) around this selection. To track the cell region in the following images the brightest pixel of the previous image is used as the midpoint to search for the new brightest and dimmest pixel as described before. Based on the fluorescence of the brightest (*F_b_*) and the dimmest (*F_d_*) pixel a threshold (*F_d_*+(*F_b_-F_d_*)/4) is defined to identify the region belonging to the cell and the background. For background correction, the intensity of the background (*F_bg_*) was subtracted from the intensity of the region of cell (*F*). The difference was divided by the mean pre-stimulus intensity of the region of cell (*F*_0_, corrected by *F_bg_*). The result is the background subtracted, relative change of the intensity of the cell region, which is reported as a percentage: *F*/*F*_0_ = 100*[(*F*-*F_bg_*)/(*F*_0_-*F_bg_*)].

The mean traces of the ASEL and the ASER dendrites were fitted individually with functions describing the ON response in ASEL and the OFF response in ASER as the difference of two exponential functions and the OFF response in ASEL and the ON response in ASER as an exponential function.
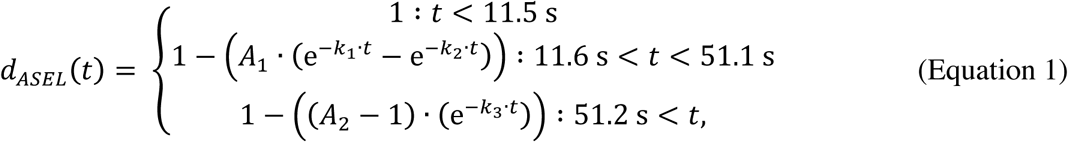

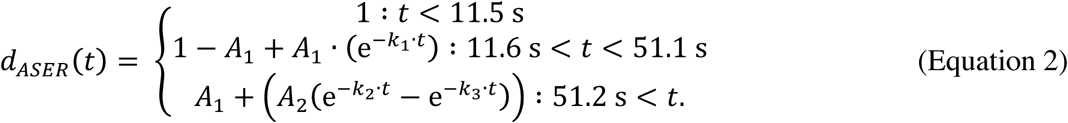

### Experimental design and statistical analyses

This study used adult *C. elegans* hermaphrodites (age: day 1, first day of egg-laying) for all experiments, which included chemotaxis assays, analysis of gene and protein expression by fluorescence microscopy, and neuronal activity by imaging a genetically-encoded calcium indicator. For each chemotaxis assay, we placed between 30-150 worms on an assay plate and analyzed at least three assay plates for each condition or condition. We pooled data from at least two independent replicates which were performed blind to genotype, except in assays involving strains carrying transgenic arrays. In the latter case, genotype was determined *post hoc* by visual inspection for the presence or absence of the transgene and behavioral performance was scored for both genotypes in parallel. To achieve this goal, we used sodium azide (0.1 M) to immobilize animals after the 30-minute chemotaxis assay period and counted animals of both genotypes on the stage of stereomicroscope equipped for epi-fluorescence (Leica M165 FC Stereomicroscope with Fluorescence).

We built and analyzed four independent transgenic lines co-expressing CMK-1::GFP from a *flp-6* promoter active in the ASE neurons and mCherry from the *cmk-1* promoter (DAG610, DAG611, DAG612, DAG613). We analyzed the subcellular position of CMK-1 using a transgenic line expressing CMK-1::GFP from a *flp-6* promoter (10-18 neurons, Figure 4A) and verified subcellular localization of strains where CMK-1 was targeted to the nucleus or cytoplasm (14-37 neurons, Figure 4C). We analyzed salt-evoked calcium transients in animals held in a microfluidic trap. In transgenic animals with the *yxEx738 [flp-6p:: GCaMP3 + unc-122p:: GFP]* in both ASE neurons, we analyzed signals in at least 10 and fewer than 27 individual ASER and ASEL neurons across conditions and genotypes. Except in rare instances (4-6 animals) we analyzed a single ASE neuron in each animal. Animals expressing *yxEx738 [flp-6p:: GCaMP3 + unc-122p:: GFP]* in either ASER or ASEL, but not both, were rare. As a result, we analyzed signals in four ASEL neurons and six ASER neurons. These data were collected during three independent imaging sessions. We generated two strains expressing *ykEx738* in a *cmk-1(ok287)* null mutant background, GN637 and GN670, and pooled the results after determining that calcium signals were indistinguishable between the two strains.

Data were tested for normality using the D’Agostino & Pearson omnibus normality test. Data passing this test were compared for differences using 2-way ANOVA followed by Sidak’s multiple comparisons test. These analyses were performed using Prism 6/7 (GraphPad). Data that did not pass normality were compared using the Wilcoxon Rank Sum test and adjusted for multiple hypothesis testing using the Benjamini-Hochberg method using R Studio. The outcomes of these statistical analyses are given in each figure legend.

## Results

As a first step in our investigation of salt aversive learning in *C. elegans*, we determined the relationship between conditioning time and the conversion of positive salt chemotaxis (attraction) to negative salt chemotaxis (aversion). We conditioned animals in buffer with or without salt in the absence of bacterial food and tested their ability to perform chemotaxis (Figure 1A). Under basal conditions, the chemotaxis index was close to +1, indicating near-perfect attraction (Materials and Methods). As shown in Figure 1B, animals converted attraction to aversion over a few hours, and the conversion followed an approximately exponential time course. This time course was well-fit by a single exponential function with a time constant of 21.7 minutes. Control animals, conditioned without salt and in the absence of bacterial food continued to perform positive chemotaxis even after 10 hours of conditioning. This analysis reveals that conditioning animals for 180 minutes (3 hours) was sufficient to saturate the aversion response. As a result, we conditioned animals for 180 minutes in the absence of bacterial food without salt (0 mM) or with 20 mM salt for all subsequent experiments unless otherwise specified.

### CMK-1 is essential for salt-learning behavior in *C. elegans*

Protein kinases are critical to rapidly transduce extracellular information about the environment to cellular and organismal responses. To understand whether and how kinases contribute to salt aversive learning, we compared the learning ability of wild-type worms to that of worms carrying mutations in several protein kinases known to be involved in relaying key environmental stimuli: AMP kinase (AMPK), Protein Kinase C (PKC), and CaM kinases (CaMKI and II). Loss-of-function mutants in the worm AMPK (*aak-2(ok524))*, PKC (*pkc-1(ok563)*, *pkc-2(ok328))*, and CaMKII genes (*unc-43(gk452)* and *unc-43(sa200))*, behaved like wild-type worms, suggesting that these genes are dispensable for both salt seeking and for salt aversive learning (Figure 1C). By contrast, *cmk-1* loss-of-function mutants, *cmk-1(ok287)* and *cmk-1(oy21)*, performed positive chemotaxis regardless of whether they were conditioned with or without salt. The defect in salt aversive learning is due to the loss of *cmk-1* function since re-expressing wild-type CMK-1 protein under its endogenous promoter was sufficient to restore salt aversive learning to *cmk-1(ok287)* mutants (Figure 1D). The defect does not appear to be due to a change in the conditioning time required for learning, however, as *cmk-1* mutants failed to convert salt attraction into aversion even after 10 hours of conditioning (CI of worms conditioned in 0 mM NaCl for 10 hours = 0.5055 +/- 0.0688 (SD), CI of worms conditioned in 20 mM NaCl for 10 hours = 0.5913 +/- 0.2056 (SD)). Note that the failure of *cmk-1* mutants to convert salt attraction into salt avoidance was not due to general defects in salt-sensing or in locomotion since naïve *cmk-1* mutants performed like wild-type animals (Figure 1B) and had average locomotion speeds similar to wild-type when assayed on a sterile agar surface (90 **μ**m/sec +/- 48 (n=37) and 115 **μ**m/s ± 55 (n=37) for *cmk-1* and wild-type, respectively). These speeds fall within the range of previously reported values (Ramot et al., 2008). Collectively, these results indicate that CaMKI, but not CaMKII or other kinases tested in this study, is essential for salt-aversive learning but not for salt chemotaxis *per se*.

### Activation of CaMKI by its upstream kinase and kinase activity is necessary for salt aversive learning

CMK-1 is a substrate of CaM kinase kinase (CaMKK, CKK-1 in worms) via phosphorylation of threonine 179 (Eto et al., 1999; Wayman et al., 2008) and *ckk-1* is co-expressed with *cmk-1* in many neurons (Kimura et al., 2002). To determine whether or not phosphorylation of CMK-1(T179) was needed for salt aversive learning, we analyzed *ckk-1* mutants and found that these mutants also exhibit defects in salt aversive learning (Figure 2A). We then tested the learning behavior of *cmk-1(ok287)* mutants overexpressing a form of CMK-1 harboring a mutation ablating the CKK-1 phosphorylation site (T179A) (Schild et al., 2014). As expected if phosphorylation of CMK-1 by CKK-1 were critical for salt aversive learning, transgenic animals expressing CMK-1(T179A) were unable to convert salt attraction into aversion (Figure 2B). Thus, *ckk-1* regulates salt aversive learning upstream of *cmk-1*, likely through phosphorylation of the CMK-1 protein.

**Figure 2.**
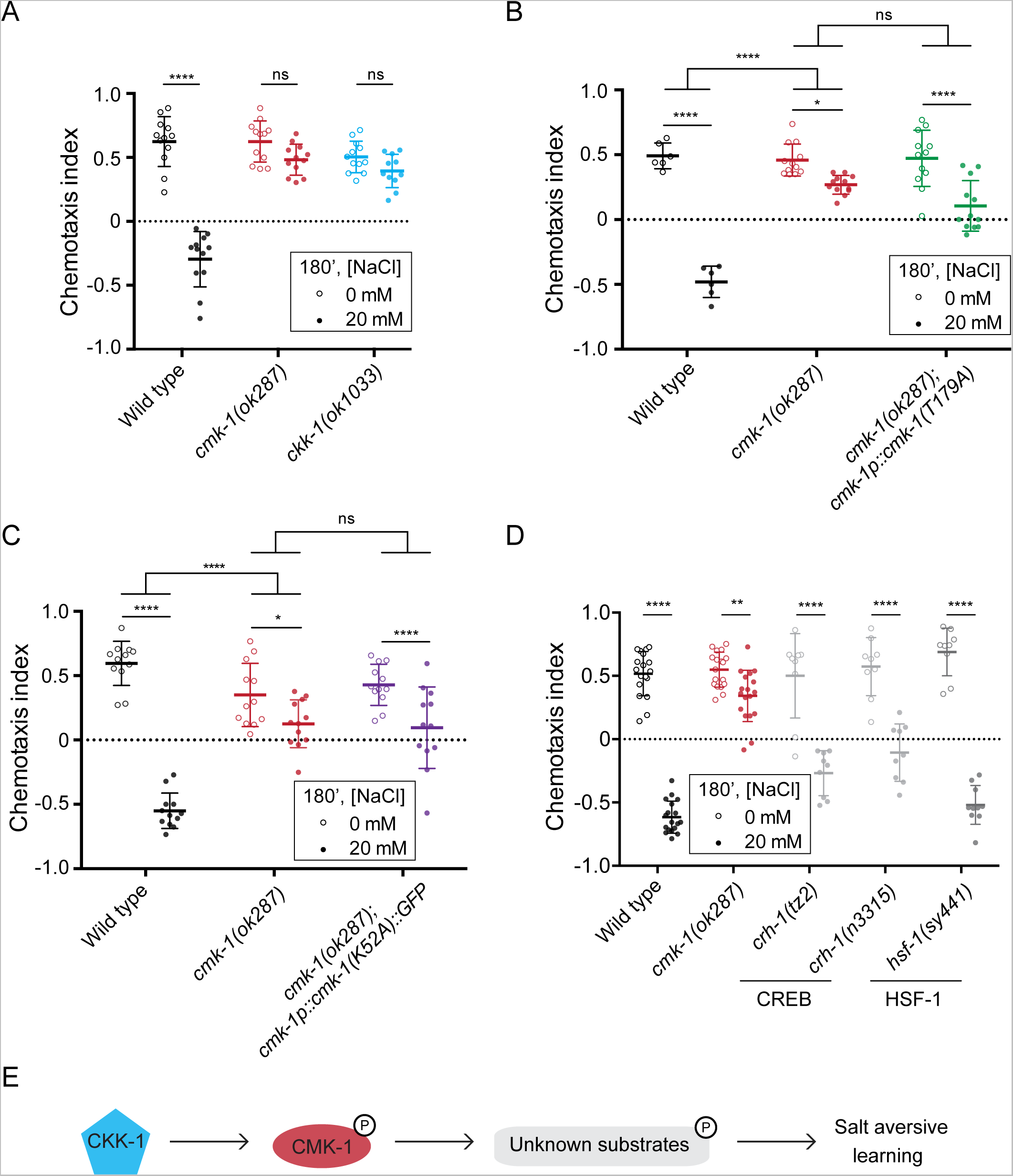
Salt aversive learning requires CaM kinase kinase, CKK-1, and CMK-1 kinase activity. **A.** CKK-1 is needed for salt aversive learning. A two-way ANOVA revealed a significant interaction between genotype and condition [*F*(2, 65) = 47.37, *p* < 0.0001] and Sidak’s multiple comparisons test revealed a significant effect of conditioning for wild type (p < 0.0001) but not *cmk-1(ok287)* or *ckk-1 (ok1033)*. **B.** A CMK-1 mutant that cannot be phosphorylated fails to restore salt aversive learning to *cmk-1* mutants. A two-way ANOVA across all genotypes and conditions revealed a significant interaction between genotype and condition [*F*(2, 54) = 26.17, *p* < 0.0001] and Sidak’s multiple comparisons test revealed a significant effect of conditioning for all genotypes. A two-way ANOVA between wild type and *cmk-1(ok287)* revealed a significant interaction between genotype and condition [*F*(1, 32) = 112, *p* < 0.0001] and a two-way ANOVA between *cmk-1(ok287)* and *cmk-1(ok287); cmk-1p:: cmk-1(T179A)* revealed a non-significant interaction between genotype and condition [*F*(1, 44) = 3.521, *p* = 0.0672]. **C.** A kinase-defective CMK-1(K52A) mutant protein cannot restore salt aversive learning to *cmk-1(ok287)* mutants. A two-way ANOVA across all genotypes and conditions revealed a significant interaction between genotype and condition [*F*(2, 66) = 33.96, *p* < 0.0001] and a Sidak’s multiple comparisons test revealed a significant effect of conditioning for all genotypes. A two-way ANOVA between wild type and *cmk-1(ok287)* revealed a significant interaction between genotype and condition [*F*(1, 44) = 71.16, *p* < 0.0001] and a two-way ANOVA between *cmk-1(ok287)* and *cmk-1(ok287); cmk-1p:: cmk-1(K52A):: GFP* revealed a non-significant interaction between genotype and condition [*F*(1, 44) = 0.6384, *p* = 0.4286]. **D.** Behavior of known CMK-1 substrates. A two-way ANOVA revealed a significant interaction between genotype and condition [*F*(4, 117) = 34.05, *p* < 0.0001]. Sidak’s multiple comparisons test was used to assess the effect of conditioning for each genotype, which was significant at a level of *p* < 0.0001 (* * * *) for all genotypes except for *cmk-1(ok287)*, which was significant at a level of *p* < 0.01 (**). For each panel in this figure, data were pooled from at least 2 independent experiments, consisting of 3-6 assays/condition performed blind to genotype. Animals of the indicated genotypes were conditioned for 180 minutes in either 0 mM or 20 mM NaCl and tested in a salt chemotaxis assay. Each symbol represents the results of a single assay; open and closed symbols indicate animals conditioned in 0 mM and 20 mM NaCl, respectively; and bars indicate mean ± SD of all assays conducted under the indicated condition and genotype. **E.** A schematic model, where CMK-1 is activated via phosphorylation by its upstream kinase CKK-1, and subsequently activates unknown downstream substrates to regulate salt aversive learning.

To determine whether the kinase activity of CMK-1 is needed for salt aversive learning, we tested a strain overexpressing CMK-1(K52A), a mutant isoform that cannot function as a kinase, but lacking endogenous CMK-1 (Schild et al., 2014). We found that transgenic worms expressing the kinase-dead CMK-1 isoform had intact salt attraction behavior, but had defects in salt aversive learning like *cmk-1(ok287)* mutants (Figure 2C). This result suggests that CMK-1 regulates salt aversive learning behavior through its function as a kinase.

CaM kinases are known to phosphorylate transcription factors such as CREB and HSF-1, thereby regulating activity-dependent changes in gene expression (Sheng et al., 1991; Eto et al., 1999; Holmberg et al., 2001; Wayman et al., 2008). However, loss-of-function mutants in CREB, *crh-1*, or HSF-1, *hsf-1*, responded to salt conditioning like wild-type animals (Figure 2D). Thus, salt aversive learning is independent of CREB and HSF-1. Similar results were observed in some (Satterlee et al., 2004; Yu et al., 2014; Kobayashi et al., 2016), but not all (Moss et al., 2016) forms of *cmk-1*-dependent plasticity studied to date. Collectively, these data implicate a pathway whereby CMK-1 is activated via phosphorylation by CKK-1 and subsequently phosphorylates downstream, yet unknown, substrates responsible for salt aversive learning (Figure 2E).

### CMK-1 acts in ASE neurons to regulate salt learning

Because CMK-1 is expressed broadly throughout the worm nervous system (Kimura et al., 2002; Satterlee et al., 2004) and proper chemotaxis behavior depends not only on salt-sensing but also on neural circuits linking sensory events to behavior, we sought to determine the neuronal locus of *cmk-1* action *vis-à-vis* salt aversive learning. Given that prior work has shown that CMK-1 acts in primary sensory neurons to regulate behavioral (Schild et al., 2014; Yu et al., 2014) and developmental (Neal et al., 2015) plasticity, we focused on the primary salt-sensing neurons, ASER and ASEL. First, we verified that CMK-1 is expressed in the ASE neurons by co-expressing CMK-1::GFP with a known ASE marker (Figure 3A). Next, we generated and tested transgenic lines that could restore wild-type CMK-1 protein to selected neurons in a *cmk-1* null mutant background. We found that expressing wild-type CMK-1 protein under the control of the ASE-specific *flp-6* promoter was sufficient to restore salt aversive learning to *cmk-1(ok287)* mutants. The ability to rescue salt aversive learning was specific to *flp-6p*, since expressing CMK-1 under the control of the mechanosensory neuron-specific *mec-3p* and the AWC chemosensory-neuron specific *ceh-36Dp* promoters failed to rescue wild-type behavior (Figure 3B). These findings indicate that not only is CMK-1 expressed in the ASE neurons, but also that CMK-1 functions in a cell-autonomous manner in these sensory neurons to control salt aversive learning behavior.

**Figure 3.**
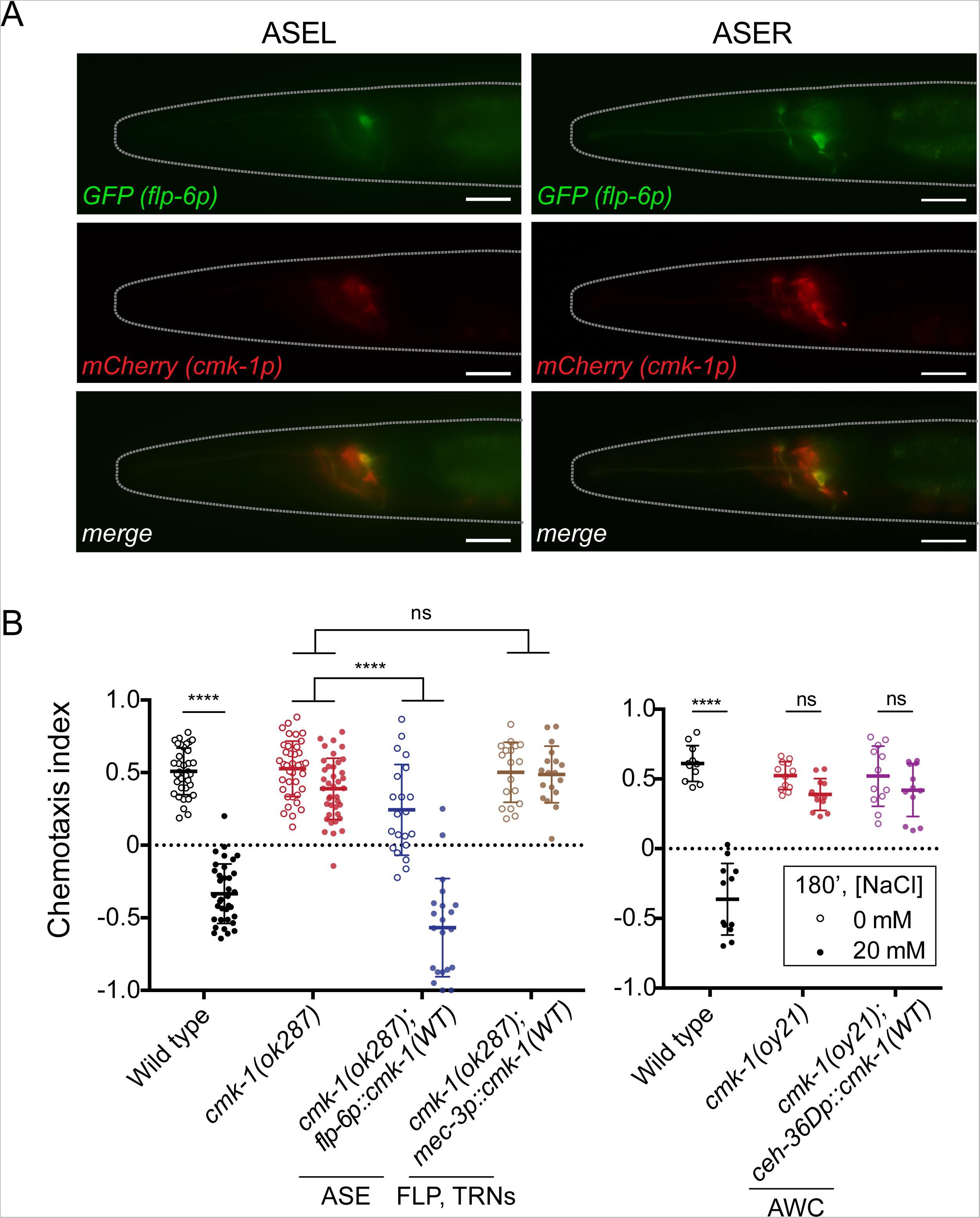
CMK-1 acts in ASE neurons to regulate learning. **A.** CMK-1 is expressed in ASE sensory neurons. GFP expressed under the *flp-6* promoter (known to drive expression in ASE neurons) co-localizes with mCherry expressed under the *cmk-1* promoter in both ASE neurons of wild type worms. Shown are the ASEL (left panels) and ASER (right panels) neurons from the same worm imaged using epifluorescence microscopy. Worms are oriented anterior side to the left and dorsal side to top of the image. Scale bar is 20 *μ*m. Similar results were observed in four (of four) independent transgenic lines. **B.** Behavior of *cmk-1(ok287)* (left) and *cmk-1(oy21)* (right) mutants expressing wild-type CMK-1 under neuron-specific promoters. Each circle represents the results of a single assay and bars indicate mean ± SD. Data were pooled from 2 independent experiments, where each experiment consisted of 3-6 plates per condition. For the left panel, a two-way ANOVA across all genotypes and conditions revealed a significant interaction between genotype and condition [*F*(3, 220) = 51, p < 0.0001] and a Sidak’s multiple comparisons test revealed a significant effect of conditioning for all genotypes except *cmk-1(ok287); mec-3p:: cmk-1(WT):: GFP*. A two-way ANOVA between *cmk-1(ok287)* and *cmk-1(ok287);flp-6p:: cmk-1(WT):: GFP* revealed a significant interaction between genotype and condition [*F*(1, 116) = 48.67, *p* < 0.0001] and a two-way ANOVA between *cmk-1(ok287)* and *cmk-1(ok287); mec-3p:: cmk-1(WT):: GFP* revealed a non-significant interaction between genotype and condition [*F*(1, 110) = 2.318, *p* = 0.1308]. For the right panel, a two-way ANOVA revealed a significant interaction between genotype and condition [*F*(2, 65) = 45.12, *p* < 0.0001] and a Sidak’s multiple comparisons test revealed a significant effect of conditioning for wild type but not *cmk-1(oy21)* or *cmk-1(oy21); ceh-36Dp:: cmk-1(WT).*

We then examined how the CMK-1 protein responded to salt stimulation in these neurons. Previous studies reported nuclear-cytoplasmic shuttling of CMK-1 in primary sensory neurons (Schild et al., 2014; Yu et al., 2014; Neal et al., 2015) and in concert with modulation of glutamate receptor expression (Moss et al., 2016). We sought to observe the subcellular localization of CMK-1 in the cell bodies of the two ASE neurons by imaging a fusion between CMK-1 and GFP. The fusion protein is sufficient to restore wild-type salt-aversive learning to *cmk-1(ok287)* mutants (Figure 1D), indicating the fusion protein functions like the wild-type protein under these conditions. Prior to conditioning, the CMK-1 protein localizes primarily to the cytoplasm of both ASE neurons (Figure 4A). We did not detect any changes in the nuclear-cytoplasmic ratio in the ASEL or ASER cell body during conditioning for between 10 and 180 minutes in 20 mM NaCl (Figure 4A). Although conditioning in salt-free buffer transiently increased the localization of CMK-1 throughout cell body in ASER neurons, the significance of this finding for salt aversive learning is unclear because learning does not occur under these conditions. We next expressed forms of CMK-1 targeted to the nucleus or the cytoplasm, thanks to the adjunction of ectopic nuclear localization signal (NLS) or nuclear export signal (NES), respectively (Figure 4B and C). In both cases, the learning defect of *cmk-1(ok287)* mutants was rescued (Figure 4D). Moreover, a mutant that encodes a form of CMK-1 with a deleted predicted NES (and a portion of the auto-inhibitory domain), *cmk-1(pg58)*, exhibited normal salt learning (Figure 4E). Collectively, these findings suggest that neither the precise localization of CMK-1 nor nuclear-cytoplasmic shuttling in the ASE cell bodies are essential for salt aversive learning.

### Calcium signals in sensory dendrites, cell bodies, and axons of ASE neurons

The two ASE neurons are bilaterally-symmetric, bipolar chemosensory neurons whose cell bodies and dendrites lie on the left and right sides of the animal and whose axons overlap in the nerve ring (Figure 5A). Despite their similar appearance, the two ASE neurons are genetically and functionally distinct: ASEL detects salt increases and ASER neurons are activated by salt decreases (Hobert, 2014). Ample evidence supports this general description of chemosensory responses in the ASE cell bodies (Tomioka et al., 2006; Suzuki et al., 2008; Ortiz et al., 2009; Thiele et al., 2009; Oda et al., 2011; Kunitomo et al., 2013; Luo et al., 2014; Rabinowitch et al., 2014), but less is known about the relationship between signals in ASE dendrites, cell bodies, and axons. We sought to fill this gap in knowledge by recording salt-evoked calcium signals in the sensory endings (dendrites), cell bodies, and axons of the wild-type ASE neurons. To reach this goal, we took advantage of mosaic expression of the *yxEx738 [flp-6p:: GCaMP3; ttx-1p:: GFP; unc-122p:: GFP]* transgene and recorded responses to pulses of sodium chloride salt in the ASEL dendrites, cell bodies, and axons independently of the ASER neuron and *vice versa*. Figure 5B shows raster plots of salt-evoked calcium signals measured in all three compartments (sensory dendrite, cell body, axon) of the ASEL and ASER neurons. Each line in the raster plot represents calcium signals as a function of time in the indicated compartment; white color indicates no change relative to the pre-stimulus baseline while red and blue colors indicate increases and decreases relative to baseline, respectively.

**Figure 4.**
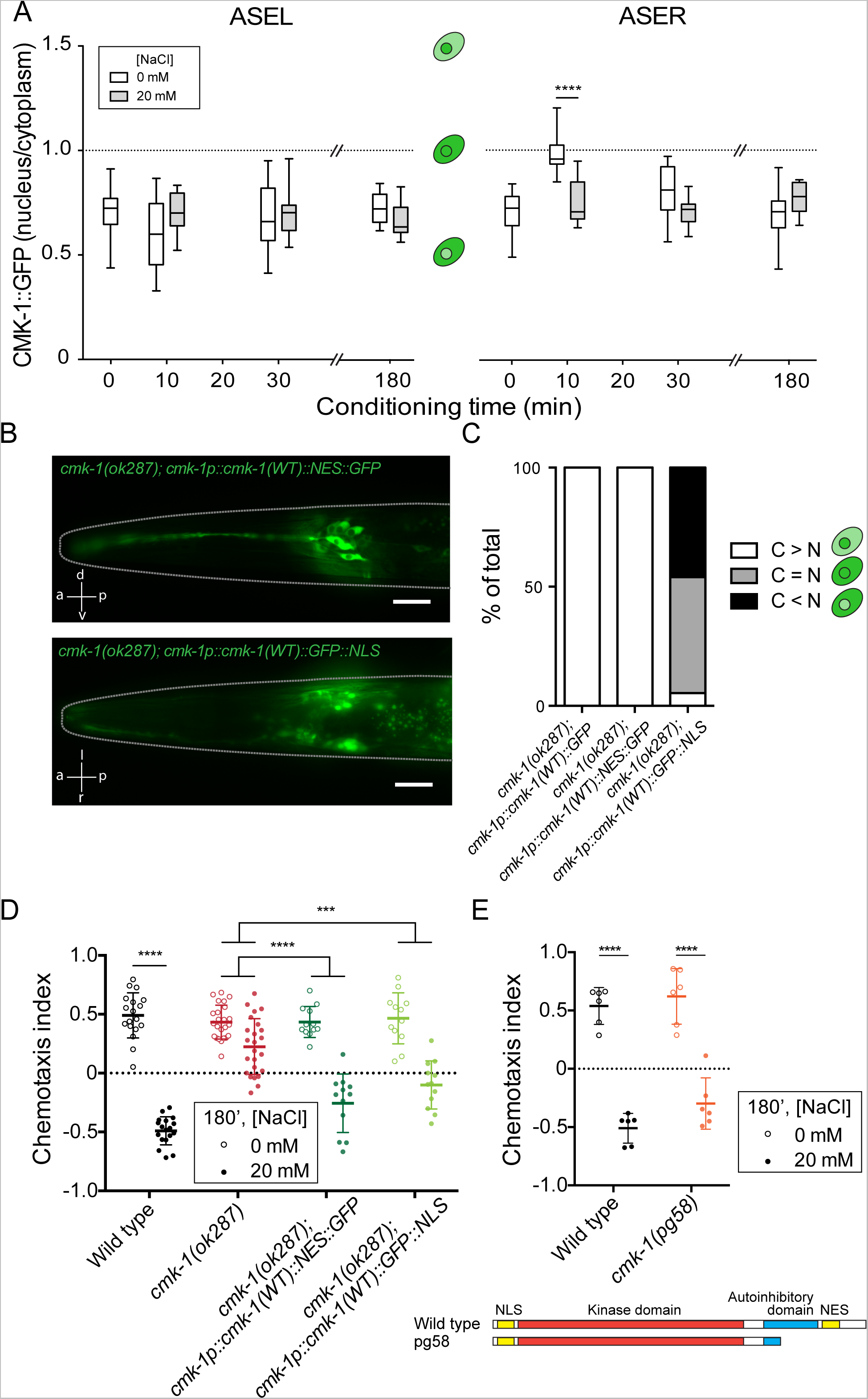
Salt conditioning does not depend on the exact subcellular localization of the CMK-1 protein. **A.** Subcellular localization of CMK-1 in the ASE neurons. The ratio of fluorescence in the nuclear compartment versus the cytoplasmic compartment is plotted separately for ASEL (left panel) and ASER (right panel). Each box-and-whisker bar is the result of the analysis of between 10-18 neurons; data were pooled from 7 independent experiments. Boxes extend from the 25^th^ to 75^th^ percentiles of the data, and whiskers extend from the minimum value to the maximum value. For ASEL, a two-way ANOVA revealed no interaction between conditioning time and condition [*F*(2, 70) = 2.346, *p* = 0.1033]. For ASER, a two-way ANOVA revealed a significant interaction between conditioning time and condition [*F*(2, 62) = 12.94, *p* < 0.0001] and a Sidak’s multiple comparisons test revealed a significant effect of conditioning at 10 min of conditioning time. **B.** Photomicrographs of transgenic animals expressing GFP-tagged CMK-1fused to an exogenous nuclear export sequence (NES) (top) or nuclear localization sequence (NLS) (bottom). The genotypes analyzed were: *cmk-1(ok287); cmk-1p::cmk-1(WT)::GFP*, *cmk-1(ok287); cmk-1p:: cmk-1(WT)::NES::GFP*, and *cmk-1(ok287); cmk-1p::cmk1(WT)::GFP::NLS*. Scale bar is 20 *μ*m. **C.** Proportion of neurons expressing more CMK-1 in the cytoplasm (C>N), the nucleus (C<N), or similar levels in both compartments (C=N). The genotypes analyzed were: *cmk-1(ok287); cmk-1p::cmk-1(WT)::GFP*, *cmk-1(ok287); cmk-1p:: cmk-1 (WT)::NES::GFP*, and *cmk-1(ok287); cmk-1p::cmk-1(WT)::GFP::NLS*. Each bar represents between 14-37 neurons. **D.** Behavior of *cmk-1(ok287)* mutants expressing CMK-1 fused to an NES or an NLS. The genotypes analyzed were: *cmk-1(ok287)*, *cmk-1(ok287); cmk-1p::cmk1 (WT)::NES::GFP*, and *cmk-1(ok287); cmk-1p::cmk-1(WT):: GFP::NLS.* Each circle is the result of a single assay and bars indicate mean ± SD. Data were pooled from at least 2 independent experiments, where each experiment consisted of 3-6 plates per condition. A two-way ANOVA across all genotypes and conditions revealed a significant interaction between genotype and condition [*F*(3, 124) = 29.21, *p* < 0.0001] and a Sidak’s multiple comparisons test revealed a significant effect of conditioning for all genotypes. A two-way ANOVA between *cmk-1(ok287)* and *cmk-1(ok287); cmk-1p:: cmk-1(WT)::NES:: GFP* revealed a significant interaction between genotype and condition [*F*(1, 68) = 23.79, *p* < 0.0001] and a two-way ANOVA between *cmk-1(ok287)* and *cmk-1(ok287); cmk-1p:: cmk1(WT):: GFP::NLS* also revealed a significant interaction between genotype and condition [*F*(1, 68) = 12.57, *p* = 0.0007]. **E.** A gain-of-function allele of *cmk-1* behaves like wild-type. Each circle represents the results of a single assay and bars indicate mean ± SD. Data were pooled from least 2 independent experiments, where each experiment consisted of 3-6 plates per condition and was blinded to genotype. A two-way ANOVA revealed a non-significant interaction between genotype and condition [F(1, 20) = 0.6954, *p* = 0.4142] and a Sidak’s multiple comparisons test revealed a significant effect of conditioning for both wild type and *cmk1 (pg58)*. The schematic (bottom) shows the protein predicted to be encoded by *cmk-1 (pg58)* and is adapted from (Schild et al., 2014).

**Figure 5.**
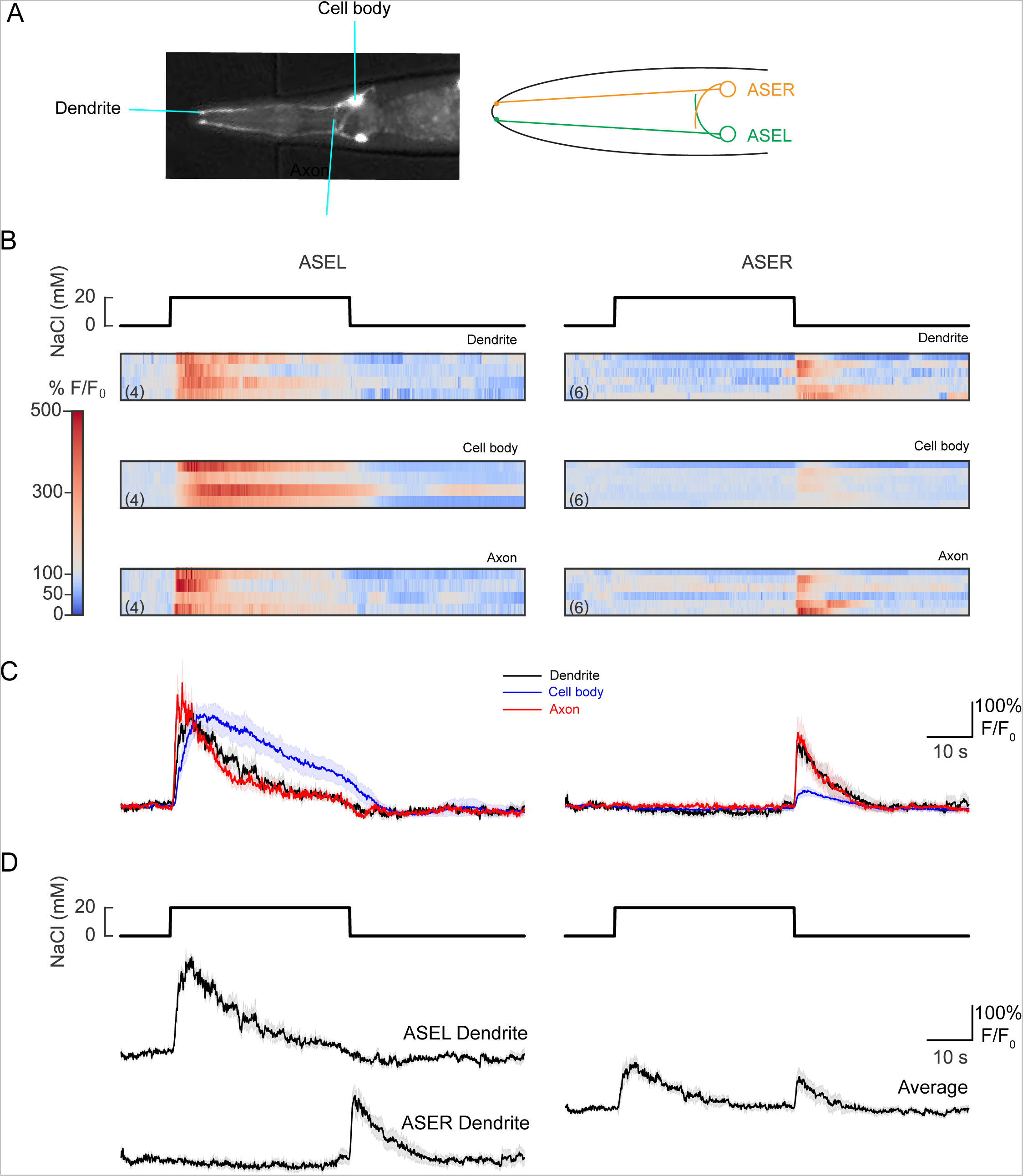
Calcium signals in wild-type ASEL and ASER dendrites, cell bodies, and axons. **A.** Image and schematic of ASE neurons. (Left) Wild type worms expressing *yxEx738 jflp-6p:: GCaMP3 + unc-122p:: GFP]*. The dendrite tips, cell bodies, and axons are easily visualized, allowing for calcium imaging of different subparts of these cells. (Right) Schematic of ASE neuron subparts. Neurite tips and cell bodies of ASEL and ASER are readily distinguishable from each other, but axons overlap in the part of the neurons that synapse onto the nerve ring of the worm. **B.** Raster plots of GCaMP3 fluorescence as a function of time in the dendrite, cell body, and axon of the ASEL (left) and ASER (right) neurons of wild type worms conditioned in 0 mM NaCl and no food and exposed to a 40 second pulse of 20 mM NaCl. Color indicates fluorescence relative to pre-stimulus control values, *F*/*F*_0_: white indicates control values, red and blue indicate increases and decreases, respectively. The number of recordings is indicated in parentheses, all of which were obtained in animals expressing *yxEx738* in ASEL or ASER, but not in both neurons. **C.** Average calcium signals in the dendrite, cell body, and axon of the ASEL (left) and ASER (right) neurons of wild type worms derived from the data shown in B. Lines indicate mean of corresponding traces and light shading indicates ± SEM. **D.** Normalized fluorescence of calcium traces in dendrites of the ASE neurons of wild type worms conditioned in 0 mM NaCl and no food and exposed to up-step (0 mM to 20 mM) and downstep (20 mM to 0 mM) of NaCl (left panel, same data as in C). Average of calcium traces in dendrites of ASEL and ASER (right panel). Lines in normalized fluorescence plots indicate mean of corresponding traces and light shading indicates ± SEM.

Salt pulses produced calcium transients synchronized to the up-step in ASEL and the down-step in ASER in all three subcellular domains (Figure 5C). Although dendritic endings and axons lie on opposite sides of the cell body and are at least 100 *μ*m apart from one another, the time course of their calcium signals were indistinguishable from one another. Responses developed more slowly and lasted longer in cell bodies than in dendrites or axons, a result that could be explained by the cell body’s larger volume acting as a low-pass filter for calcium signals. These observations are compatible with the electrically compact nature of the ASE neurons (Goodman et al., 1998) and a lack of distinct, subcellular calcium domains in these cells (Kuramochi and Doi, 2017).

We averaged the mean responses of ASEL and ASER axons and used this to predict the waveform expected in animals expressing a calcium indicator in both axons (Figure 5D). These reconstructed traces recapitulate the phasic responses we detected in animals expressing *yxEx738* in both ASE axons (Figure 7E, 7F). With this analysis, we confirm that ASEL functions as an ON cell that is sensitive primarily to salt up-steps whereas ASER functions as an OFF cell that responds primarily to salt down-steps. We establish that signals recorded in sensory dendrites, but not cell bodies, mirror the stimulus-induced calcium waveforms present in axons. We further propose that signals recorded from the nerve ring of animals expressing calcium indicator in both neurons can be interpreted as the average of the axonal signals generated independently by ASEL and ASER.

### Loss of *cmk-1* blunts the effect of salt conditioning on somatic calcium signals

Calcium signals recorded in the cell bodies of specific neurons are a familiar correlate of learning behaviors (Clark et al., 2006; Biron et al., 2008; Yu et al., 2014). Thus, to understand the impact of *cmk-1* mutation on calcium signals, we compared chemosensory responses in ASE cell bodies of wild type and *cmk-1* mutants in response to salt conditioning (Figure 6). In wild-type worms, salt conditioning sufficient to transform behavioral attraction to aversion *decreased* ON responses in ASEL (Figure 6A, 6B, 6E, left) by approximately 2-fold. Similar effects have been reported in animals conditioned in 20 mM NaCl for only 10 minutes (Oda et al., 2011), suggesting that plasticity in chemosensory signaling can develop rapidly. Calcium levels were suppressed during salt application in ASER cell bodies following conditioning in the presence of 20 mM NaCl (Figure 6A - B, right).

**Figure 6.**
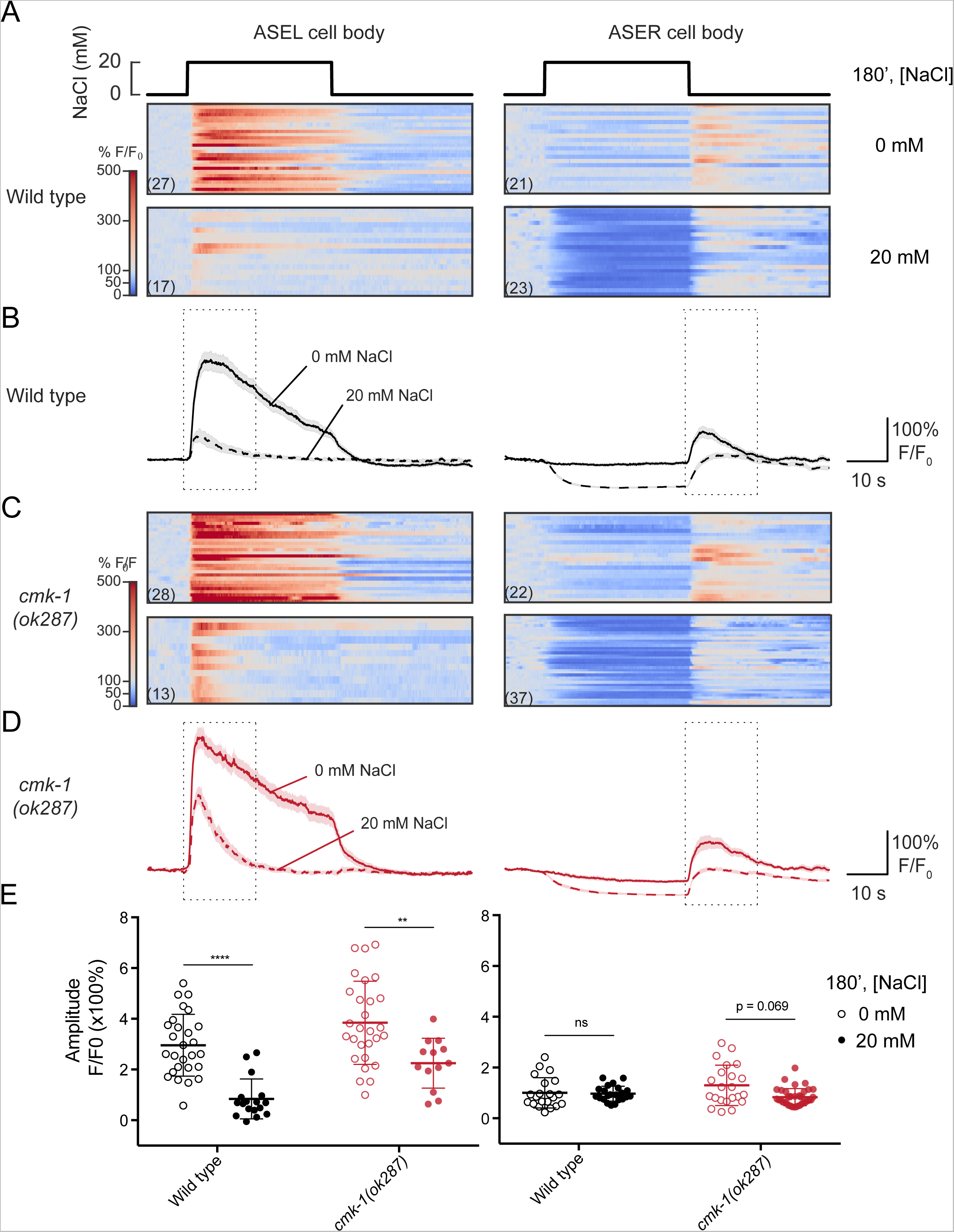
The effect to salt conditioning and *cmk-1* loss of function on calcium signals in ASER and ASEL cell bodies. **A.** Raster plots of GCaMP3 fluorescence as a function of time in the in cell bodies of the ASEL (left) and ASER (right) neurons of wild type worms conditioned in 0 mM or 20 mM NaCl and no food and exposed to a 40 second pulse of 20 mM NaCl. Each horizontal line corresponds to a recording from one cell body; the number of recordings is indicated in parentheses. Color indicates fluorescence relative to pre-stimulus control values, *F*/*F*_0_: white indicates control values, red and blue indicate increases and decreases, respectively. **B.** Average calcium signals in the cell bodies of the ASEL (left) and ASER (right) neurons of wild type worms. The traces are derived from the data shown in A. Lines indicate the mean and light shading indicates ± SEM. **C.** Raster plots of GCaMP3 fluorescence as a function of time in the cell bodies of the ASEL (left) and ASER (right) of *cmk-1(ok287)* worms conditioned in 0 mM or 20 mM NaCl and no food and exposed to a 40 second pulse of 20 mM NaCl. Each horizontal line corresponds to a recording of a single cell body; the number of recordings is indicated in parentheses. Color indicates fluorescence relative to pre-stimulus control values, *F*/*F*_0_: white indicates control values, red and blue indicate increases and decreases, respectively. **D.** Average fluorescence of calcium traces in cell bodies of the ASEL (left) and ASER (right) neurons of *cmk-1(ok287)* worms conditioned in 0 mM or 20 mM NaCl and no food and exposed to a 40 second pulse of 20 mM NaCl. The traces are derived from the data shown in C. Lines indicate the mean and light shading indicates ± SEM. **E.** Maximum amplitude of the change in calcium signals recorded in cell bodies of the ASEL (left) and ASER (right) neurons of wild type and *cmk-1(ok287)* worms conditioned in 0 mM or 20 mM NaCl and no food. The amplitude was extracted from calcium traces by subtracting the average signal of 10 s before the outlined region from the maximum signal in the region outlined by dotted line in the normalized fluorescence plots in panels B and D. Data were analyzed using the Wilcoxon Rank Sum test for non-parametric data, where p-values were adjusted for multiple hypothesis testing using the BenjaminiHochberg method. For ASEL cell body, 20 mM NaCl conditioned cell bodies of both wild type and *cmk-1(ok287)* exhibited significantly lower maximum fluorescence intensities upon upshift from 0 to 20 mM NaCl compared with 0 mM NaCl conditioned cells (wild type *p* < 0.0001, *cmk-1(ok287) p* < 0.01). For ASER cell body, 20 mM NaCl conditioned cell bodies of neither wild type nor *cmk-1(ok287)* ASEL exhibited significantly lower maximum fluorescence intensities upon downshift from 0 to 20 mM NaCl compared with 0 mM NaCl conditioned cells (wild type *p* = 0.5728, *cmk-1(ok287) p* = 0.0691).

**Figure 7.**
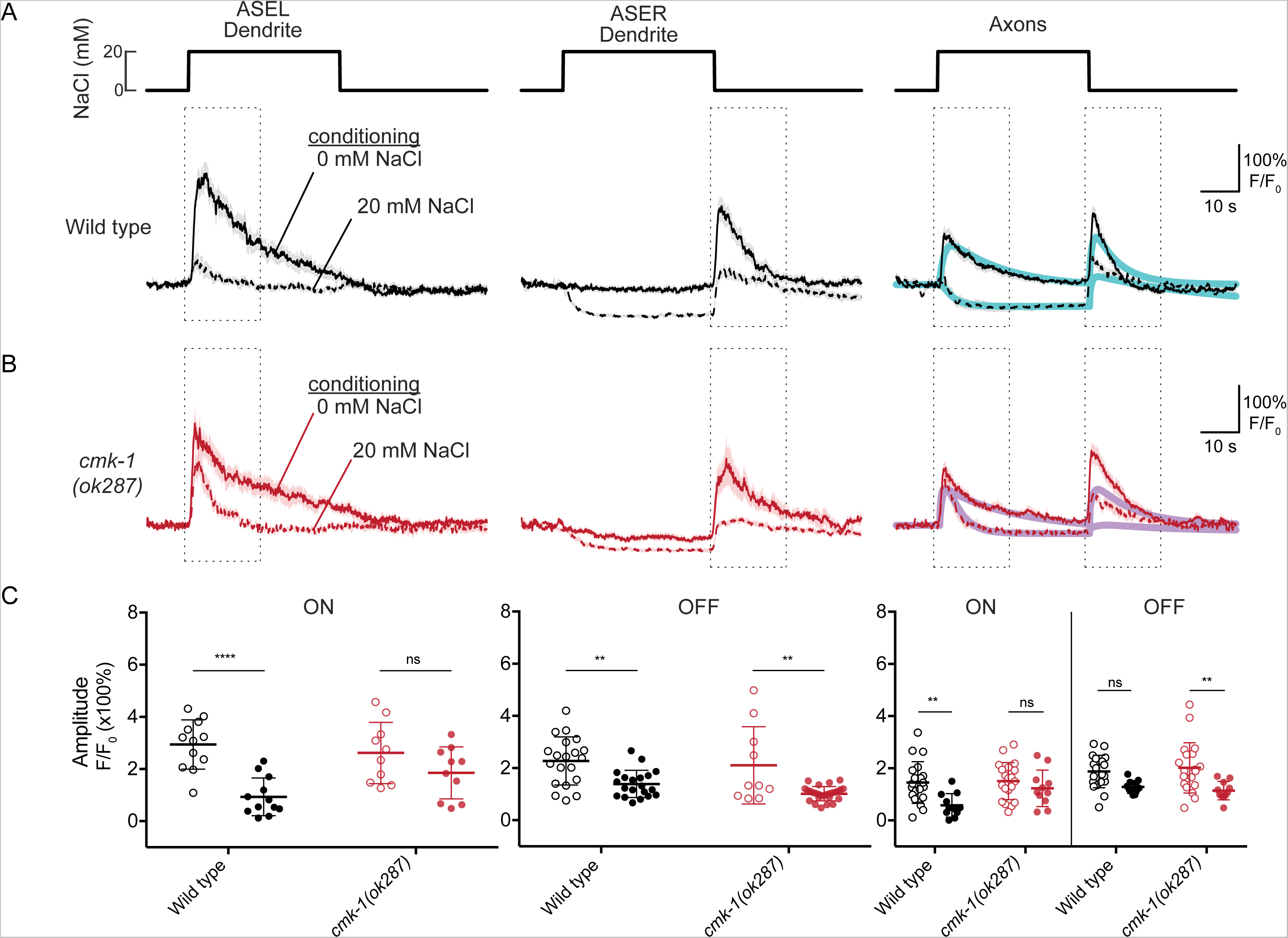
CMK-1 is required for the effect of salt conditioning on chemosensory calcium signals in dendrites and axons of ASE sensory neurons. **A.** Average chemosensory calcium signals in the dendrites of wild-type ASEL (left) and ASER (right) neurons of wild type worms conditioned in 0 mM or 20 mM NaCl and no food and exposed to a 40 second pulse of 20 mM NaCl. Lines indicate the mean and light shading indicates ± SEM of 12 and 20 recordings in ASEL and ASER dendrites, respectively. Consult Extended Data Panel A for the source data. **B.** Average chemosensory calcium signals recorded from ASEL (left) and ASER (right) dendrite tips of *cmk-1(ok287)* worms conditioned in 0 mM or 20 mM NaCl and no food and exposed to a 40 second pulse of 20 mM NaCl. Lines indicate the mean and light shading indicates ± SEM of 10, 10, 20, and 27 recordings from ASEL dendrites conditioned without or with 20 mM NaCl and ASER dendrites conditioned without or with 20 mM NaCl, respectively. Consult Extended Data Panel B for the source data. **C.** Maximum amplitude of ON (left) and OFF (right) responses in dendrites of ASE neurons of wild type and *cmk-1(ok287)* worms conditioned in 0 mM NaCl (open circles) or 20 mM NaCl (closed circles). Maximum amplitude was extracted from calcium traces by subtracting the average signal of 10 s before the outlined region from the maximum signal in the region outlined by dotted line in the normalized fluorescence plots in Fig 7A and 7B. For the ASEL dendrite, a two-way ANOVA revealed a significant interaction between genotype and condition [*F*(1, 40) = 4.546, *p* = 0.0392] and Sidak’s multiple comparisons test revealed a significant effect of conditioning for wild type (adjusted *p* < 0.000 1), but not *cmk-1(ok287)* (adjusted *p* = 0.4042). For the ASER dendrite, a two-way ANOVA failed to detect an interaction between genotype and condition [*F*(1, 73) = 0.303, *p* = 0.5837] and a Sidak’s multiple comparisons test revealed a significant effect of conditioning for both wild type (adjusted *p* = 0.0031) and *cmk-1(ok287)* (adjusted *p* = 0.0016). **D.** Average chemosensory calcium signals in the axons of ASE neurons in wild type worms conditioned in 0 mM or 20 mM NaCl and no food and exposed to a 40 second pulse of 20 mM NaCl. Lines indicate the mean and light shading indicates ± SEM of 20 and 11 recordings in animals conditioned without and with 20 mM NaCl, respectively. Consult Extended Data Panel C for the source data. The solid cyan lines are a weighted average of the mean traces of the ASEL and ASER dendrites, which were fitted individually (*d_ASEL_*(*t*), *d_ASER_*(*t*); fitting function described in methods section). The weighting factor, *d_Axon_*(*t*)=*A*\**d_ASEL_*(*t*)+(1-*A*)\**d_ASER_*(*t*)_,_ was estimated by minimizing the r^2^of the difference between *d_Axon_*(*t*) and the mean fluorescence signal measured from the axon (black line). The weighting factors were 0.4 and 0.29 for 0 mM and 20 mM NaCl conditioned animals. **E.** Average chemosensory calcium signals in the axons of ASE neurons in *cmk-1(ok287)* worms conditioned in 0 mM or 20 mM NaCl and no food and exposed to a 40 second pulse of 20 mM NaCl. Lines indicate the mean and light shading indicates ± SEM of 20 and 11 recordings in animals conditioned without and with 20 mM NaCl, respectively. Consult Extended Data Panel D for the source data. The solid magenta lines are a weighted average of fits to the mean traces of the ASEL and ASER dendrites (*d_ASEL_*, *d_ASER_*; fitting function described in methods section). The weighting factor, *d_Axon_*=*A*^*^*d_ASEL_*+(1-*A*)^*^*d_ASER_* by the r^2^, was estimated minimizing of the difference between *d_Axon_* and the mean fluorescence signal measured from the axon. The weighting factor was 0.47 and 0.67 for 0 mM and 20 mM NaCl conditioned animals, respectively. **F.** Maximum amplitude of ON (left) and OFF (right) responses in the axons of ASE neurons of wild type and *cmk-1(ok287)* worms conditioned in 0 mM NaCl (open circles) or 20 mM NaCl (closed circles). Maximum amplitude was extracted from calcium traces by subtracting the average signal of 10 s before the outlined region from the maximum signal in the region outlined by dotted line in the normalized fluorescence plots in Fig 7D and 7E. A two-way ANOVA failed to reveal an interaction between genotype and condition [*F*(3, 116) = 1.203, *p* = 0.3119] and a Sidak’s multiple comparisons test revealed a significant effect of conditioning for the ON response in wild type (adjusted p = 0.0042) and the OFF response in *cmk-1(ok287)* (adjusted *p* = 0.0040), but not for ON responses in *cmk-1(ok287)* (adjusted *p* = 0.7427) or the OFF response in wild type (adjusted *p* = 0.0970).

Interestingly, *cmk-1(ok287)* mutants exhibited qualitatively similar changes in calcium responses in the cell bodies of both ASEL and ASER upon conditioning with 20 mM NaCl compared to wild type (Figure 6C-D). The conditioning-dependent decreases in ON responses in ASEL were blunted in *cmk-1* mutants compared to wild type (Figure 6E). These results indicate that ASEL and ASER neurons in *cmk-1(ok287)* mutants retain their function as ON and OFF cells, respectively, but differ from wild-type in their response to conditioning. Although it remains unclear how changes in ASE signals govern behavioral plasticity, these observations are consistent with our conclusion that *cmk-1* acts in the ASE neurons to support salt aversive learning.

### *cmk-1* mutants exhibit defects in calcium signals in ASER and ASEL dendrite tips and axons

Given that the response dynamics in ASE cell bodies are slowed compared to signals observed at the tip of the ASE dendrites and along their axons (Figure 5), we also determined the effect of salt conditioning and loss of *cmk-1* function on calcium signals recorded in these subcellular domains. Consistent with current and prior findings in the cell bodies, ASEL sensory endings detect up-shifts in salt and ASER endings detect downshifts. In wild type worms, the peak amplitude of the ON response in ASEL dendrites and of the OFF response in ASER dendrites decreased after salt conditioning (Figure 7A, Extended Data A, left and center). This effect of conditioning was diminished in *cmk-1* mutant ASEL dendrites (Figure 7B, Extended Data B, left and center). We observed a qualitatively similar, but less dramatic effect in *cmk-1* mutant ASER dendrites. We summarized these results by comparing peak calcium values observed at the onset (ASEL) and offset (ASER) of the salt pulse (Figure 7C). A two-way ANOVA revealed a significant effect of conditioning on the amplitude of ON responses in ASEL [*F*(1, 40) = 22.67, p < 0.0001] and interaction between genotype and conditioning in ASEL [*F*(1, 40) = 4.546, p = 0.0392], but only a significant effect of conditioning on the amplitude of OFF responses in ASER [*F*(1, 73) = 27.72, p < 0.0001]. Because the dendrite tips are the site of chemosensory transduction, these results suggest that *cmk-1* is required for conditioning-dependent chemosensory plasticity or adaptation.

Finally, we analyzed how conditioning in 20 mM NaCl and loss of *cmk-1* function affected responses in the overlapping ASE axons (Figure 7D – F), which ultimately transmit changes in sensory neuron activity to downstream interneurons and subsequently to motor neurons to affect the link between sensation and behavior. Conditioning wild type animals in 20 mM NaCl had two main effects on axonal calcium signals in ASE neurons. First, responses to salt up-steps are transformed from a transient increase in axonal calcium to a sustained decrease in calcium (Figure 7D, Extended Data A, right panel). Second, OFF responses to salt down-steps decreased slightly (Figure 7D, Extended Data A, right panel). Conditioning in 20 mM NaCl failed to generate analogous changes in *cmk-1* mutant axons (Figure 7E, Extended Data B, right panel), however. In particular, *cmk-1* mutant ASE axons failed to generate wild-type-like conditioned responses to salt up-steps and down-steps (Figure 7E - F, Extended Data B, right panel).

We considered these observations in the context of our earlier findings that: 1) chemosensory signals in overlapping axons are consistent with a summation of the signals generated by the ASEL and ASER axons (Figure 5D) and that 2) the chemosensory waveforms recorded in single ASE dendrites are virtually indistinguishable from those recorded in axons (Figure 5C). Assuming these relationships hold for *cmk-1* mutant ASE neurons, we fit chemosensory waveforms measured in wild type and *cmk-1* ASER and ASEL dendrites (Materials and Methods), computed a weighted sum of these waveforms and compared the resulting waveforms to the measured axonal signals. As shown in Figure 7D (cyan curves), this simple strategy was sufficient to recapitulate the effects of conditioning by 20 mM NaCl on chemosensory signaling in wild type animals, reaffirming that conditioning depresses calcium levels during salt pulses. Although this approach could reproduce the responses to salt up-steps in *cmk-1* mutants, it was less successful at reproducing the responses to salt down-steps (Figure 7E, purple curves).

Compatible with the fact that *cmk-1* mutants perform chemotaxis like wild-type animals when conditioned with 0 mM NaCl (Figure 1C), these imaging results establish that *cmk-1(ok287)* mutant ASE neurons retain stimulus-evoked calcium signals similar to those found in wild-type neurons. In contrast, the response of *cmk-1(ok287)* mutant ASE neurons diverged from wild-type upon salt conditioning at the site of sensory transduction and synaptic communication. Whereas calcium responses in wild-type animals decrease in ASEL and ASER, we detected little if any change in the size of ON responses in *cmk-1(ok287)* mutants. Collectively, our data show that the loss of salt aversive learning ability in *cmk-1* mutants correlates with specific alterations in the calcium transients at multiple locations in ASE neurons.

## Discussion

CMK-1, the *C. elegans* ortholog of mammalian CaMKI/IV, enables behavioral plasticity related to thermal acclimation and in developmental plasticity governed by chemosensation (Satterlee et al., 2004; Schild et al., 2014; Yu et al., 2014; Neal et al., 2015). Here, we demonstrate that CMK-1 is also critical for salt aversive learning, a process that allows animals to balance the need for salt with the potentially damaging effects of this essential inorganic nutrient. We show that salt aversive learning depends on CMK-1 expression in the ASE chemsensory neurons, employs an upstream activation pathway involving CKK-1, and requires the kinase activity of CMK-1.

Consistent with prior work, we show that ASEL functions as an ON cell and ASER functions as an OFF cell and that salt conditioning blunts salt-evoked calcium signals. We analyzed stimulus-evoked calcium signals in the dendrites, cell bodies, and axons of both ASER and ASEL neurons, extending general knowledge of chemosensory signaling by these neurons. We found that signals in dendrites and axons were indistinguishable from one another and that somatic signals resembled a low-pass filtered version of these signals. These findings are consistent with isopotential electrical signaling by ASE (Goodman et al., 1998) and have implications for relying on somatic calcium signals as a proxy for global activity in the *C. elegans* nervous system.

Salt-evoked calcium signals in *cmk-1* mutant ASE neurons are less sensitive to salt conditioning than they are in their wild-type counterparts. We propose that this defect in sensory signals accounts for the inability of *cmk-1* mutants to perform salt aversive learning. The fact that we detected differences in the response to conditioning in the ASEL and ASER dendrites implies that conditioning acts *via* a *cmk-1*-dependent pathway to alter chemosensory transduction. ASE signaling is linked to behavior through the ASE axon, which contains both presynaptic and postsynaptic sites. To understand more about the effect of salt conditioning on signaling by wild-type and *cmk-1* axons, we leveraged our finding that signal dynamics were similar in wild-type dendrites and axons. Specifically, we hypothesized that signal dynamics in the overlapping axons could be predicted from a weighted sum of the signals in ASEL and ASER dendrites. This simple model recapitulated the principle features of the phasic responses to salt pulses we observed in wild-type ASE axons, but was not adequate to capture the effect of salt conditioning in *cmk-1* mutant axons (Figure 7E). Thus, CMK-1 is likely to play additional roles in ASE signal transmission in addition to its effect on the plasticity of chemosensory transduction.

CMK-1 is expressed globally in the *C. elegans* nervous system (Kimura et al., 2002; Satterlee et al., 2004), but has been shown to act locally in sensory neurons to support plasticity (Schild et al., 2014; Yu et al., 2014; Neal et al., 2015). Its kinase activity is required for plasticity, but CREB, a canonical CaM kinase substrate, is not essential for the forms of behavioral and developmental plasticity analyzed thus far. Although we cannot exclude the possibility that CREB acts redundantly or that it is not involved in these forms of learning, it seems likely that CMK-1 relies on other substrates and that the relevant substrates might be neuron-specific. Important next steps will be to discover additional proteins phosphorylated by CMK-1 and to analyze their contribution to behavioral plasticity related to chemosensory and thermosensory plasticity.

In conclusion, we have characterized the ability of a single gene, *cmk-1*, to profoundly impact learning in *C. elegans*. This, along with the sufficiency of this gene in a single pair of neurons to rescue this behavior, illustrates a relatively simple circuit in this animal to adapt to the natural world. The fact that worm CMK-1 is thought to serve functions performed by two different proteins in mammals, CaMKII and CaMKI/CaMKIV (Cohen et al., 2015), speaks further to the efficiency of this system. Evolutionary advantages of such a model include the ability to perform relatively complex behaviors with very few neurons and molecular players. Potential risks, however, may include insufficient compensatory mechanisms to rescue the system when damaged. The ability to learn to avoid areas of high salt upon starvation likely evolved as a way for organisms to minimize energy exploring barren areas and instead seek food-rich habitats.

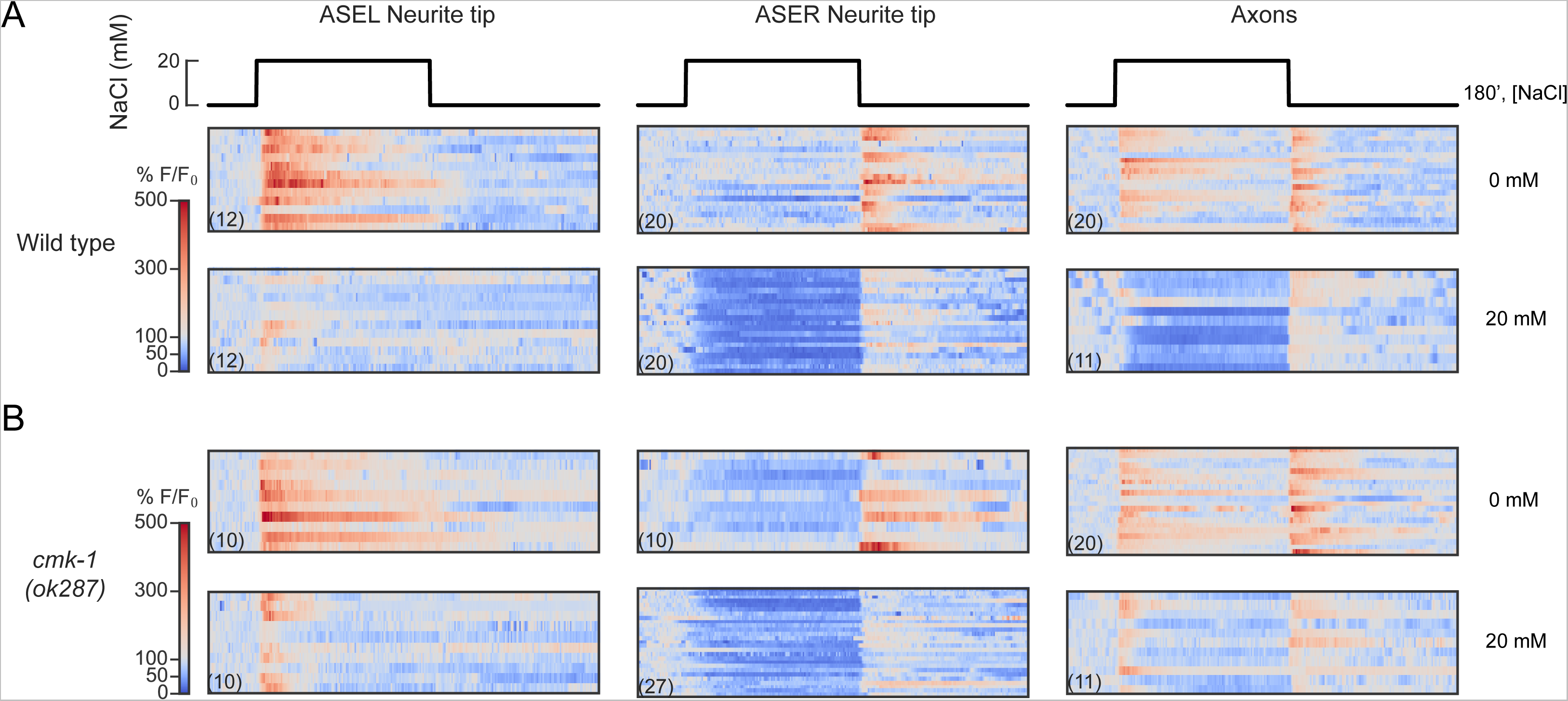

## Acknowledgements

We thank C. Murphy (Princeton), Y. Zhang (Harvard), and the Caenorhabditis Genetics Center, which is funded by NIH Office of Research Infrastructure Programs (P40 OD010440), for worm strains, and S. Chalasani (Scripps) for technical advice regarding microfluidics set-up. We also thank our colleagues Z. Liao, V. Lauper, and L. Schild for superb technical support, and J. Kubanek for help with worm tracking. We are grateful to L. Booth, B. Dulken, S. Han, C-K. Hu, K. Papsdorf, and P. Singh for critical reading of the manuscript. This research was funded by a National Science Foundation Graduate Research Fellowship (J.P.L.), NIH grant NS047715 (M.B.G), NIH grant AG031198 (A.B.), and Swiss National Science Foundation grants BSSGI0_155764 and PP00P3_150681 (D.A.G.).

## Extended Data

**A.** Raster plots of chemosensory calcium signals in the ASEL and ASER dendrites in wild type worms conditioned in 0 mM or 20 mM NaCl and no food and exposed to a 40 second pulse of 20mM NaCl. Each horizontal line corresponds to a recording from a single dendrite tip; the number of recordings is shown in parentheses. Color indicates the change in fluorescence relative to pre-stimulus levels: red indicates an increase and blue indicates a decrease.

**B.** Raster plots of chemosensory calcium signals in the ASEL and ASER dendrites in *cmk-1(ok287)* mutants conditioned in 0 mM or 20 mM NaCl and no food and exposed to a 40 second pulse of 20 mM NaCl. Each horizontal line corresponds to a recording from a single dendrite tip; the number of recordings is shown in parentheses. Color indicates the change in fluorescence relative to pre-stimulus levels: red indicates an increase and blue indicates a decrease.

**C.** Raster plots of chemosensory calcium signals in the axons of ASE neurons in wild type worms conditioned in 0 mM or 20 mM NaCl and no food and exposed to a 40 second pulse of 20 mM NaCl. Each horizontal line corresponds to a recording from a single dendrite tip; the number of recordings is shown in parentheses. Color indicates the change in fluorescence relative to pre-stimulus levels: red indicates an increase and blue indicates a decrease.

**D.** Raster plots of chemosensory calcium signals in the axons of ASE neurons in *cmk-1(ok287)* mutants conditioned in 0 mM or 20 mM NaCl and no food and exposed to a 40 second pulse of 20 mM NaCl. Each horizontal line corresponds to a recording from a single dendrite tip; the number of recordings is shown in parentheses. Color indicates the change in fluorescence relative to pre-stimulus levels: red indicates an increase and blue indicates a decrease.

